# Development and worldwide use of a non-lethal and minimal population-level impact protocols for the isolation of chytrids from amphibians

**DOI:** 10.1101/246538

**Authors:** Matthew C. Fisher, Pria Ghosh, Jennifer M. G. Shelton, Kieran Bates, Lola Brookes, Claudia Wierzbicki, Gonçalo M. Rosa, Rhys A. Farrer, David M. Aanensen, Mario Alvarado-Rybak, Arnaud Bataille, Lee Berger, Susanne Böll, Jaime Bosch, France C. Clare, Elodie Courtois, Angelica Crottini, Andrew A. Cunningham, Thomas M. Doherty-Bone, Fikirte Gebresenbet, David J. Gower, Jacob Höglund, Timothy Y. James, Thomas S. Jenkinson, Tiffany A. Kosch, Carolina Lambertini, Anssi Laurila, Chun-Fu Lin, Adeline Loyau, An Martel, Sara Meurling, Claude Miaud, Pete Minting, Serge Ndriantsoa, Frank Pasmans, Tsanta Rakotonanahary, Falitiana C. E. Rabemananjara, Luisa P. Ribeiro, Dirk S. Schmeller, Benedikt R. Schmidt, Lee Skerratt, Freya Smith, Claudio Soto-Azat, Giulia Tessa, Luís Felipe Toledo, Andrés Valenzuela-Sánchez, Ruhan Verster, Judit Vörös, Bruce Waldman, Rebecca J. Webb, Che Weldon, Emma Wombwell, Kelly R. Zamudio, Joyce E. Longcore, Trenton W.J. Garner

**Affiliations:** Department of Infectious Disease Epidemiology, School of Public Health, Faculty of Medicine (St Mary’s campus), Imperial College London, London W2 1PG, UK; Centro de Investigación para la Sustentabilidad, Facultad de Ecología y Recursos Naturales, Universidad Andres Bello, Republica 440, Santiago, Chile; Laboratory of Behavioral and Population Ecology, School of Biological Sciences, Seoul National University, Seoul 08826, South Korea; UMR ASTRE, CIRAD, Montpellier, France; One Health Research Group, College of Public Health, Medical and Veterinary Sciences, James Cook University, Townsville, Queensland, 4811, Australia; Agency for Population Ecology and Nature Conservancy, Gerbrunn, Germany; Museo Nacional de Ciencias Naturales, CSIC c/ Jose Gutierrez Abascal 2, 28006 Madrid, Spain; Laboratoire Ecologie, évolution, interactions des systèmes amazoniens (LEEISA), Université de Guyane, CNRS, IFREMER, 97300 Cayenne, French Guiana.; CIBIO - Centro de Investigação em Biodiversidade e Recursos Genéticos, InBIO, Universidade do Porto, 4485-661 Vairão Portugal; Institute of Zoology, Regent’s Park, London, NW1 4RY, UK; Conservation Programmes, Royal Zoological Society of Scotland, Edinburgh, UK; Department of Integrative Biology, Oklahoma State University, 113 Life Sciences West, Stillwater, OK 74078, USA; Life Sciences, The Natural History Museum, London SW7 5BD, UK; Department of Ecology and Genetics, EBC, Uppsala University. Norbyv. 18D, SE-75236, Uppsala, Sweden; Zoology Division, Endemic Species Research Institute, 1 Ming-shen East Road, Jiji, Nantou 552, Taiwan; Helmholtz Centre for Environmental Research – UFZ, Department of Conservation Biology, Permoserstrasse 15, 04318 Leipzig, Germany; ECOLAB, Université de Toulouse, CNRS, INPT, UPS, Toulouse, France; Department of Pathology, Bacteriology and Avian Diseases, Faculty of Veterinary Medicine, Ghent University, Salisburylaan 133, B-9820 Merelbeke, Belgium; PSL Research University, CEFE UMR 5175, CNRS, Université de Montpellier, Université Paul-Valéry Montpellier, EPHE, Biogéographie et Ecologie des vertébrés, Montpellier, France; Amphibian and Reptile Conservation (ARC) Trust, 655A Christchurch Road, Boscombe, Bournemouth, Dorset, UK, BH1 4AP; Durrell Wildlife Conservation Trust, Madagascar Programme, Madagascar; Department of Evolutionary Biology and Environmental Studies, University of Zurich, Winterthurerstrasse 190, 8057 Zurich, Switzerland; National Wildlife Management Centre, APHA, Woodchester Park, Gloucestershire GL10 3UJ, UK; Non-profit Association Zirichiltaggi - Sardinia Wildlife Conservation, Strada Vicinale Filigheddu 62/C, I-07100 Sassari, Italy; ONG Ranita de Darwin, Nataniel Cox 152, Santiago, Chile; Unit for Environmental Sciences and Management, Private Bag x6001, North-West University, Potchefstroom, 2520, South Africa; Collection of Amphibians and Reptiles, Department of Zoology, Hungarian Natural History Museum, Budapest, Baross u. 13., 1088, Hungary; School of Biology and Ecology, University of Maine, Orono, Maine 04469, USA; Department of Ecology and Evolutionary Biology, University of Michigan, Ann Arbor, Michigan, 48109, USA; Laboratório de História Natural de Anfíbios Brasileiros (LaHNAB), Departamento de Biologia Animal, Instituto de Biologia, Universidade Estadual de Campinas, Campinas, São Paulo, 13083-862, Brazil; Department of Ecology and Evolutionary Biology, Cornell University, Ithaca, New York, 14853, USA; Centre for Ecology, Evolution and Environmental Changes (CE3C), Faculdade de Ciências da Universidade de Lisboa, Lisboa, Portugal; IUCN SSC Amphibian Specialist Group-Madagascar, 101 Antananarivo, Madagascar; Info Fauna Karch, Université de Neuchâtel, Bellevaux 51, UniMail Bâtiment 6, 2000 Neuchâtel, Switzerland; Centre for Genomic Pathogen Surveillance, Wellcome Genome Campus, Cambridgeshire, UK

## Abstract

Parasitic chytrid fungi have emerged as a significant threat to amphibian species worldwide, necessitating the development of techniques to isolate these pathogens into sterile culture for research purposes. However, early methods of isolating chytrids from their hosts relied on killing amphibians. We modified a pre-existing protocol for isolating chytrids from infected animals to use toe clips and biopsies from toe webbing rather than euthanizing hosts, and distributed the protocol to interested researchers worldwide as part of the BiodivERsA project *RACE* – here called the *RML* protocol. In tandem, we developed a lethal procedure for isolating chytrids from tadpole mouthparts. Reviewing a database of use a decade after their inception, we find that these methods have been widely applied across at least 5 continents, 23 countries and in 62 amphibian species, and have been successfully used to isolate chytrids in remote field locations. Isolation of chytrids by the non-lethal *RML* protocol occured in 18% of attempts with 207 fungal isolates and three species of chytrid being recovered. Isolation of chytrids from tadpoles occured in 43% of attempts with 334 fungal isolates of one species (*Batrachochytrium dendrobatidis*) being recovered. Together, these methods have resulted in a significant reduction and refinement of our use of threatened amphibian species and have improved our ability to work with this important group of emerging fungal pathogens.

## INTRODUCTION

A major consequence of globalisation has been the increase of invasive species owing to trade in live animals and plants. A further outcome of this process is the concomitant rise of novel emerging fungal pathogens (EFPs; (Farrer *et al*. 2017)) as these infections are moved within trade networks and establish in uninfected regions – an example of fungal ‘pathogen pollution’ (Fisher *et al*. 2012). Whilst EFPs can affect humans, they have also been broadly detrimental to natural populations of plants and animals, leading to worldwide losses of biodiversity. This dynamic has been most apparent across amphibians, where EFPs leading to population extirpation and species extinctions have contributed to amphibians now being the most endangered class of vertebrate (Stuart *et al*. 2004; Mendelson *et al*. 2006). In particular, emergence of parasitic fungi in the genus *Batrachochytrium* (phylum Chytridiomycota, order Rhizophydiales) have played a major role in driving amphibian population and species declines worldwide (Berger *et al*. 1998; Fisher *et al*. 2009).

While a single species, *Batrachochytrium dendrobatidis* (*Bd*), was originally thought to have caused the ongoing panzootic (James *et al*. 2009), we now know that amphibian chytridiomycosis is caused by a much broader swathe of phylogenetic diversity than was previously thought (Farrer *et al*. 2011; Schloegel *et al*. 2012). Next-generation sequencing and phylogenomic analyses have shown that *Bd sensu stricto* is composed of deep genetic lineages which are emerging through international trade in amphibians (Fisher *et al*. 2007; Schloegel *et al*. 2009; Schloegel *et al*. 2010). Superimposed upon this background of trade-associated lineages of *Bd* has come the recent discovery of a new species of pathogenic chytrid, also within the Rhizophydiales, *B. salamandrivorans* (*Bsal*; Martel *et al*. 2013). This pathogen has rapidly extirpated European fire salamanders (*Salamandra salamandra*) in the Netherlands and a broad screening of urodeles has shown that *Bsal* occurs naturally in southeast Asia where it appears to asymptomatically infect salamander and newt species (Laking *et al*. 2017).

The ability to isolate and culture both *Bd* and *Bsal* has played a key role in catalysing research into their pathogenesis and virulence (Voyles *et al*. 2007; Rosenblum *et al*. 2012; Farrer *et al*. 2017), phenotypic characteristics (Piotrowski *et al*. 2004; Fisher *et al*. 2009; Becker *et al*. 2017) and a wealth of experimental studies on epidemiologically relevant parameters (Garner *et al*. 2009; Ribas *et al*. 2009; Rosenblum *et al*. 2012). Longcore *et al*. (1999) first isolated *Bd* from infected amphibians by modifying techniques used to isolate other chytrids (Barr 1987). Longcore cleaned small (< 0.5mm dia) pieces of *Bd*-infected leg and foot skin by wiping them through agar and then placed skin pieces onto a clean plate of nutrient agar containing penicillin G and streptomycin. This method worked well for isolating from dead animals sent by courier from North and Central America. The method, however, requires euthanizing potentially healthy animals if their infection status was unknown. Further, it is difficult to perform this protocol in remote regions that lack suitable laboratory facilities, and the lethal sampling of amphibians may be contraindicated if the species is endangered, protected or located in protected areas.

We confronted this issue in a 2008-2014 project funded by BiodivERsA (<u>http://www.biodiversa.org</u>) – *RACE*: Risk Assessment of Chytridiomycosis to European amphibian biodiversity (Fisher *et al*. 2012). One of the objectives of this project was to adjust the protocol of Longcore *et al*. (1999) to (i) reduce the need to kill adult amphibians, (ii) improve rates of chytrid isolation by allowing the use of more animals, (iii) develop protocols that enabled isolation in a field setting, and, (iv) integrate the data into the GPS-smartphone enabled epidemiological software application *Epicollect* (Aanensen *et al*. 2009; Aanensen *et al*. 2014). Further, ‘forewarned is forearmed’ and we wished to determine whether the protocol was able to isolate other species of chytrid that are part of the amphibian skin microbiota, and that may present a biosecurity risk. This need to more broadly characterise global chytrid biodiversity was met by using resources from *RACE* to train researchers worldwide in chytrid isolation techniques to provide opportunities to characterise novel chytrids as they were discovered.

In addition to the non-lethal isolation protocol, a lethal method was developed in parallel to isolate chytrids from the mouthparts of larval amphibians. We describe this method as a refinement to the main isolation protocol.

## METHODS

### Non-lethal field isolation of chytrids

Animals were captured and held in separate plastic bags or suitable containers until ready for processing (Supp. Info. *RML* Protocol 1 and Supp. Info. Swabbing Protocol 2). Using clean gloves and sterilized dissection scissors or scalpel blades, the terminal 1-2mm of the phalanges of the 4^th^ hind toe (counting from the proximal toe) was clipped and laid on the surface of an mTGhL + antibiotic (200 mg/L penicillin-G and 400 mg/L streptomycin sulphate) agar plate. Alternatively, ~1mm toe-webbing biopsy punches were taken (Sklar instruments, PA, USA) then laid on a plate. This allowed multiple animals to be processed rapidly in the field. Subsequently, each tissue sample was transferred to a second plate with a sterile needle or forceps then cleaned (as far as possible) of surface-contaminating bacteria and fungi by dragging it through the agar-medium. The needle or forceps was then used to place the tissue sample in a sterile 2 ml screw-cap microtube containing liquid mTGhL medium with antibiotics, then stored in a cool, dry place. While 4 °C appears optimal, we have successfully used shaded regions of streams to cool cultures when refrigeration was not immediately available and have even held tubes and plates for several days at > 10 °C until suitable storage conditions were available.

Once back in the laboratory, samples in tubes were visually screened for evidence of yeast or bacterial contamination (when the media takes on a ‘cloudy’ appearance), or mycelial ‘balls’ around the toe that are evidence of non-chytrid fungal contaminants. Visibly clear samples were decanted into a single well of a sterile 12-well lidded culture plate then incubated at 18°C for up to 4 weeks, topping up with extra medium to counter evaporation as necessary. Depending on the size of the initial tissue sample, toe clips and webbing were divided into several smaller samples before transferring to liquid culture media.

### Isolating chytrids from tadpoles

Tadpoles often have higher burdens of infection than adults, especially long-lived tadpoles (Skerratt *et al*. 2008), and have higher densities and encounter rates than adults. In some situations where tadpoles were large and infections heavy, tadpoles were microscopically screened with a dissecting microscope or hand lens for areas of dekeratinization of the mouth parts, especially the jaw sheaths, that indicates infection (Fellers *et al*. 2001; Smith *et al*. 2007). Tadpoles are killed before excising their mouthparts and these preliminary microscopic screens enabled us to use only a small number of animals to isolate chytrids. Additionally, uninfected and naïve tadpoles that were reared in captivity were used as live substrates to bait chytrids from adult amphibians with low levels of *Bd* infection (Bataille *et al*. 2013).

Susceptible tadpoles were reared until gills were resorbed and animals were free-swimming and feeding (developmental Gosner stage 25), because at earlier stages they are still developing the keratinized mouthparts. Each tadpole container was then immersed within a similar but larger container that held at least one chytrid-infected animal. Water exchange between the infected and bait animal containers occurred through small holes (< 0.3 mm) drilled into the bottom of the walls of the smaller internal containers. Animals were held in these conditions for between 2 and 4 weeks at species-appropriate conditions. Tadpoles were periodically examined every fourth day for the presence of the depigmented areas in the jaw sheaths that have been associated with chytrid infection.

Isolating chytrids from tadpoles first required killing by immersion in a 5 g/L solution of MS-222 (Torreilles *et al*. 2009) or other approved method. Note that anaesthetics which contain ethanol, such as phenoxyethanol (Gentz 2007), should be avoided as these will kill chytrids while MS222 is not toxic (Webb *et al*. 2005). We then dissected out keratinized jaw sheaths and cleaned the entire sheath, or sections, as above using an agar plate with antibiotics ((Longcore *et al*. 1999); Supp. Info. *RML* Protocol 1). Cleaned sections were then placed singly into sterile 12-well culture plates with 1 mL liquid media + antibiotics, or onto agar plates with 6 – 10 sections per plate, and incubated at 10 – 20 °C.

Because zoospore release may occur immediately, especially from tadpole mouthparts, cultures were examined with an inverted microscope for the presence of active zoospores every day for up to one week following the day that they were initiated. After that, checks every two days were sufficient.

### Culture and diagnosis of chytrid isolates

Subsequent culture methods for *Bd* followed those of Longcore *et al*. (1999). When isolation of *Bsal* was anticipated an incubation temperature of 15 °C was required (Blooi *et al*. 2015) whereas a temperature of 18 – 22 °C is closer to the measured growth optimum of *Bd* (Longcore *et al*. 1999; Ribas *et al*. 2009). Once growth of zoospores and/or zoosporangia was observed, 100 – 500 µL volume of culture containing zoospores and zoosporangia was transferred by pipette to a new 12-well plate with liquid medium and no antibiotics, and incubated at 15 – 20 °C. All successfully cultured isolates were subcultured into larger volumes, then centrifuged at 1700 g for 10 min before cryopreservation. A portion of the initial pellet was also be used for DNA extraction, while the remaining volume was resuspended in 10% DMSO and 10% FCS in liquid media and transferred into six 2 mL cryotubes for cryopreservation at -80 °C (Boyle *et al*. 2003).

We confirmed the identity of *Bd* and *Bsal* by quantitative PCR with an MGB Taqman probe assay in either single-plex or multiplex (Boyle *et al*. 2004; Blooi *et al*. 2013). We identified non-*Batrachochytrium* chytrids was achieved by sequencing appropriate regions of the ribosomal RNA gene with universal fungal primers followed by comparison against OTUs held in UNITE database (Unified system for DNA-based fungal species linked to classification: <u>https://unite.ut.ee</u>) to establish a species-hypothesis for the chytrid isolate in question (Schoch *et al*. 2012). If further genetic data were required, then multilocus analysis or whole-genome sequencing was undertaken using chytrid-specific methods (James *et al*. 2009; Farrer *et al*. 2013; Farrer *et al*. 2017; Farrer *et al*. 2017).

### Collation of data

To track and report chytrid isolation for the *RACE* project, we used a generic data collection tool that allows the collection and submission of geotagged data forms from field locations, *Epicollect5* (<u>https://five.epicollect.net</u>). This software has the advantage that it can be used on mobile devices with or without internet connection, and allows the immediate sharing of data across the research community. Our database at <u>https://five.epicollect.net/project/bd-global-isolation-protocol</u> included the following data fields: Date; Continent, Country, Site name; Latitude/Longitude; Wild caught or trade?; Amphibian species; Life history stage; Number sampled; Chytrid isolated?; Number isolated; Species of chytrid isolated; Chytrid lineage; Photograph of amphibian; Name of researchers.

## RESULTS

The ‘*RACE* modified Longcore (*RML*) Protocol’ for the non-lethal isolation of chytrids from amphibians is detailed in Supp. Info. 1. Ensure that you have the relevant licences, permits and permissions from ethical committees to follow the *RML* protocol 1, swabbing protocol 2 and isolation from larval amphibians.

### Non-lethal isolation from adult and juvenile amphibians

Following the formalisation and distribution of the *RACE* protocols, our Epicollect5 project summarised chytrid surveys from 2007 through to 2017 (Table 1). The Epicollect5 database can be spatially visualised at <u>https://five.epicollect.net/project/bd-global-isolation-protocol/data</u>. Figure 1 depicts the isolation of amphibian-associated chytrids using the *RACE* protocols from 5 continents (Africa, Asia, Australia, Europe and South America), 23 countries, 239 sampling episodes, and from latitudes spanning -44.1 S (*Batrachyla antartandica*, Chile) through to 55.6 N (*Bufo viridis,* Sweden). Chytrids have been non-lethally isolated from 34 amphibian species, of which 28 were anuran and 5 were caudatan species. The database also contains 5 records of chytrids that were non-lethally sampled from the amphibian trade.

**TABLE 1.** Non-lethal isolation of chytrids from adult and juvenile amphibians

| Continent | Country | *n* Species<sup>1</sup> | *n* Sampled<sup>2</sup> | *n* Chytrid<sup>3</sup> | Chytrid species |
| --- | --- | --- | --- | --- | --- |
| Africa | Madagascar | 2 | 145 | 2 | *Kappamyces* sp. |
|  | Cameroon | 1 | 30 | 1 | *B. dendrobatidis* |
|  | Ethiopia | 1 | 5 | 1 | *B. dendrobatidis* |
|  | South Africa | 6 | 179 | 45 | *B. dendrobatidis* |
| Asia | South Korea | 2 | 28 | 10 | *B. dendrobatidis* |
|  | Taiwan | 3 | 103 | 13 | *B. dendrobatidis*/<br>*Kappamyces* sp. |
| Australia | Australia | 1 | 2 | 2 | *B. dendrobatidis* |
| Europe | Belgium | 1 | 11 | 2 | *B. dendrobatidis* |
|  | France | 2 | 261 | 70 | *B. dendrobatidis* |
|  | Hungary | 1 | 15 | 3 | *B. dendrobatidis* |
|  | Italy | 1 | 14 | 4 | *B. dendrobatidis* |
|  | Portugal | 1 | 5 | 1 | *Rhizophydium*<br>sp. |
|  | Spain | 4 | 198 | 37 | *B. dendrobatidis* |
|  | Sweden | 1 | 23 | 5 | *B. dendrobatidis* |
|  | Switzerland | 1 | 30 | 1 | *B. dendrobatidis* |
|  | UK | 4 | 50 | 8 | *B. dendrobatidis* |
| South America | Chile | 1 | 10 | 1 | *B. dendrobatidis* |
|  | French Guiana | 2 | 66 | 2 | *B. dendrobatidis* |
| Trade | n/a | 4 | 15 | 5 | *B. dendrobatidis* |

<sup>1</sup>Number of amphibian species sampled, <sup>2</sup>total numbers of amphibians sampled, <sup>3</sup>number of chytrids isolated

**Figure 1.**
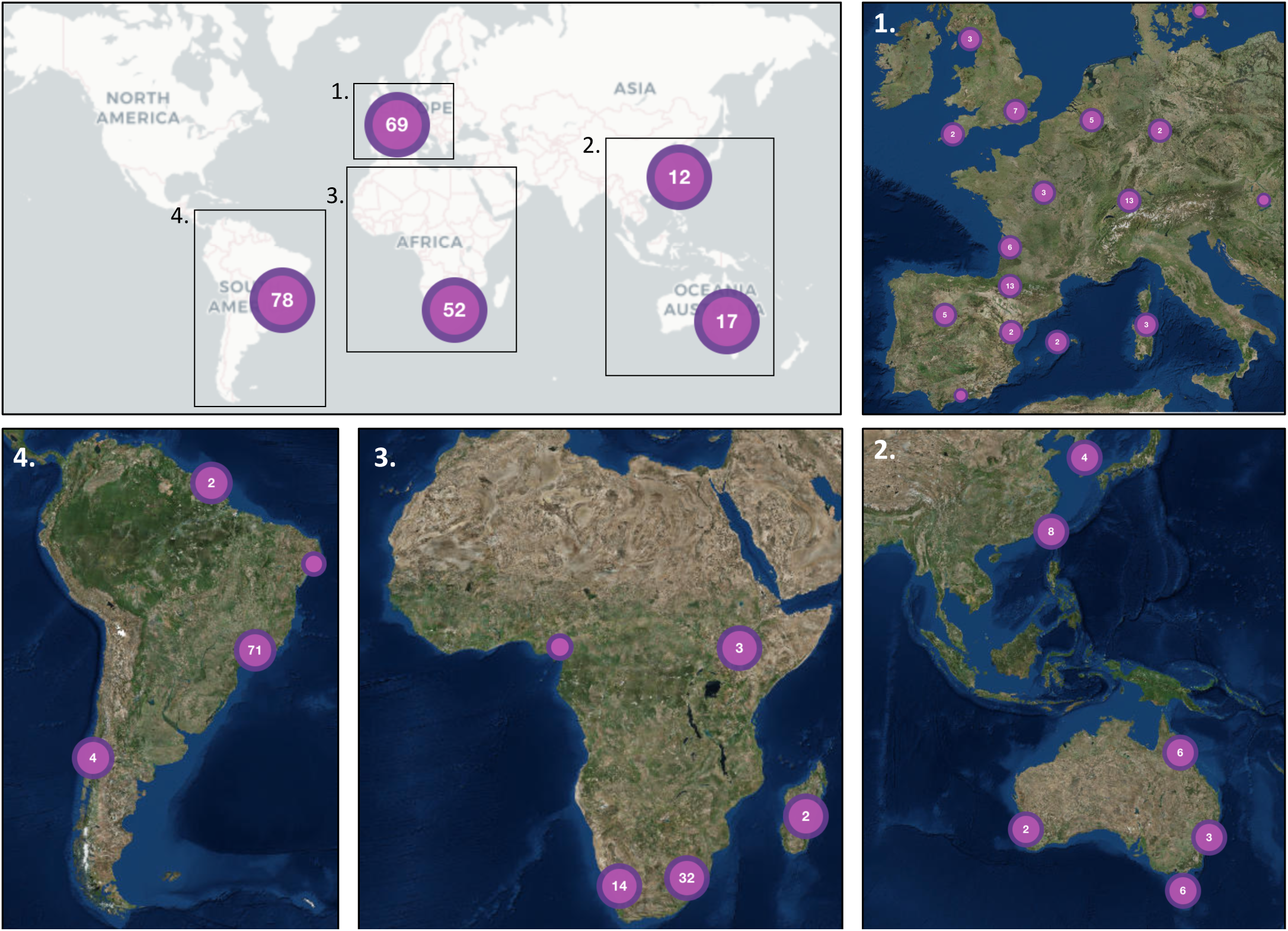
Worldwide distribution of sites where the *RML* Longcore protocol has been used to isolate chytrids. Numbers denote the quantity of amphibian species investigated. A browseable version of this *Epicollect 5* map can be accessed at https://five.epicollect.net/project/bd-global-isolation-protocol

In total, 1,152 animals were non-lethally sampled, recovering 207 chytrid isolates and resulting in a recovery rate of 18% (~1 isolate per 5 animals sampled). Of these chytrids, 203 (98%) were *Bd*, 2 were *Rhizophydium* sp., 2 were *Kappamyces* sp. and none were *Bsal* (Table 1). Of the *Bd* isolated, 42 (88%) were determined to be *Bd*GPL, 5 (10%) were *Bd*CAPE, and 1 (2%) was *Bd*CH.

### Isolation of chytrids from larval amphibians

In total, 784 tadpoles were sampled recovering 334 chytrid isolates and resulting in a recovery rate of 43% (~1 isolate per 2 – 3 animals sampled). Isolates were recovered from 34 species of amphibian, all of which were anurans (Table 2). These chytrid isolates were all *Bd* and, of the lineages recorded, 129 (78%) were *Bd*GPL, 34 (20%) were *Bd*BRAZIL and 3 (2%) were hybrids.

Baiting chytrid isolates from live adult animals using tadpoles was used successfully in South Korean *Bombina orientalis* as previously described (Bataille *et al*. 2013). Here, six tadpoles were co-housed with adult *B. orientalis*, yielding a single isolate of *Bd* for each attempt equating to a rate of success of ~20%.

**TABLE 2.**
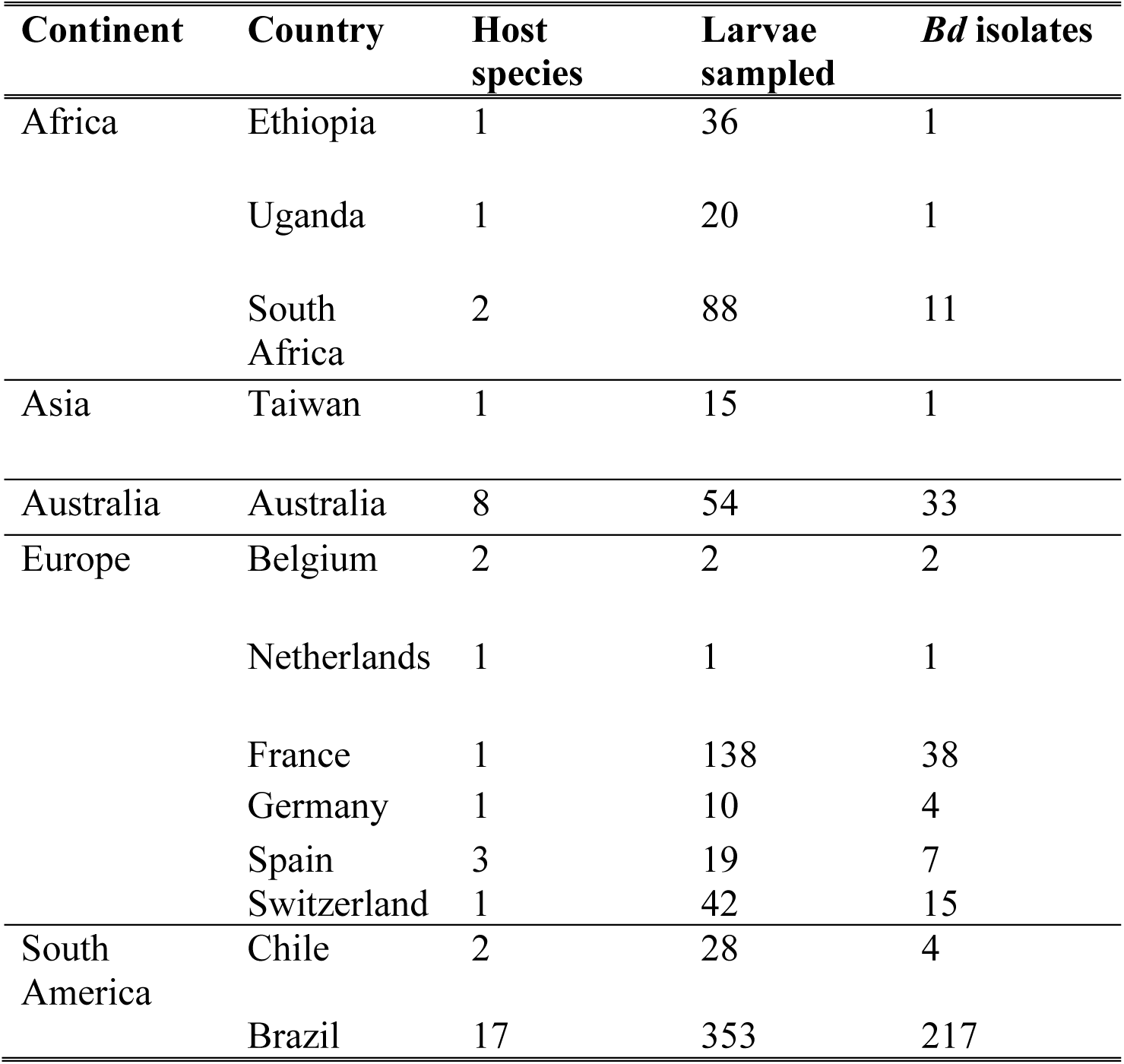
Isolation of *Batrachochytrium dendrobatidis* from mouthparts of larval amphibians

## DISCUSSION

The *RML* protocol, based on the original suggestions of Joyce Longcore for the non-lethal isolation of chytrids from amphibians, has been a success with isolates of chytrids recorded from five continents. There are likely many other unrecorded uses of this method because this protocol has been widely dispersed during the 5-year span (2008-2014) of the *RACE* project which trained a cohort of amphibian disease researchers in these techniques.

In some circumstances chytrids could not be recovered from toe-clips when sampling populations with persistent infection despite repeated attempts. This was particularly evident when the prevalence and burden of chytrid infections in surveys was low (Swei *et al*. 2011; Bataille *et al*. 2013; Laking *et al*. 2017) or when host species occupied habitats with high bacterial and/or non-target fungal contaminants. In these situations we isolated chytrids from tadpole mouthparts as an associated method to the *RML* protocol. The value of the *RML* protocol in propelling forward research on amphibian chytridiomycosis has been very clear: for instance, of the 59 scientific papers produced by *RACE*, 15 directly used isolates of *Bd* that were generated by this protocol for experimental trials (Supp. Info. 3). Further, subsequently many more studies using these isolates have extended our knowledge of the genetic diversity of *Bd* (James *et al*. 2009; Farrer *et al*. 2011; Farrer *et al*. 2013; Jenkinson *et al*. 2016), the development of novel diagnostics (Dillon *et al*. 2017), the genetic repertoire that underpins the virulence of these pathogens (Rosenblum *et al*. 2012; Farrer *et al*. 2017) and the biogeographic distributions of *Bd* diversity worldwide (Farrer *et al*. 2011; Jenkinson *et al*. 2016).

Clearly some uncontrolled biases and unanswered questions in these studies need attention. First, the majority of *Bd* isolates belong to the *Bd*GPL lineage. This could be because this lineage is more widespread (and therefore more readily recovered) than other lineages (James *et al*. 2015), or it could be that the intensity of *Bd*GPL infections and/or the rate of zoospore production is higher than for other lineages, which would also equate to a higher rate of isolation. To achieve a true and unbiased understanding of the distribution of these lineages, a lineage-specific diagnostic will need to be developed and deployed. Second, if lineage-specific differences in the probability of successful isolation exist, then mixed infections where these lineages co-occur may not be detected. This can be controlled for by isolating and genotyping many isolates from a single host and population, although this may not fully account for this bias. A related bias is that not all infectious species of chytrid will respond equally to culturing attempts. For instance, despite known attempts to isolate *Bsal* from across its endemic southeast Asian range using the protocol, to date no successful isolations of *Bsal* have been recorded. This is likely due to a combination of the low prevalence and burden of infection in salamanders and newts combined with the low initial growth-rate of *Bsal* (Martel *et al*. 2013; Laking *et al*. 2017). With the *RML* protocol, however, workers have been able to isolate non-*Bd* species of chytrid (*e.g., Kappamyces* spp. and *Rhizophydium* sp. Table 1). This diversity likely represents only a fraction of the diversity of amphibian-associated chytrids that occur, and non-biased estimators of this diversity by, for instance, profiling the nuclear ribosomal RNA cistron (Schoch *et al*. 2012), are sorely needed.

In this age of the global amphibian crisis, research on the effects of chytrid infections is transitioning to attempts to mitigate their impacts (Schmeller *et al*. 2014; Garner *et al*. 2016; Canessa *et al*. 2018). Both of these research streams benefit from the availability of chytrid isolates, but the ethics behind these research programs can be improved. To that end, our data on isolation success suggest that tadpoles are a better target for isolation than metamorphosed animals. This is to some degree unfortunate, because isolation from tadpoles requires killing. However we have outlined one refinement where captive reared tadpoles can be used to ‘bait’ infections from wild-caught amphibians to isolate chytrids without killing adult amphibians. Here, it is important to recognise that amphibians which have been co-housed in collections should not be returned to the wild due to the danger of cross-transmission of pathogens during husbandry (Walker *et al*. 2008). If it is necessary to isolate chytrids directly from wild tadpoles without using bait animals, we suggest that researchers focus on more fecund species with long larval periods as the focal species in aquatic amphibian communities. Removal of small numbers of tadpoles when clutch sizes are in the hundreds or thousands means that removals will have an insignificant ecological impact; for this reason sacrificing tadpoles is preferable to killing adult animals.

The extent to which toe-clipping effects the fitness of amphibians has been much debated (*e.g.* May (2004) but see Funk *et al*. (2005)). Toe-clipping has been shown to decrease amphibian survival, but this effect, when present, is linearly related to the number of toes removed (McCarthy *et al*. 2004; Ulmar Grafe *et al*. 2011). For the single toe-clip that the *RML* protocol requires, reduction in survival appears to be negligible (Ott *et al*. 1999; Funk *et al*. 2005), and toe clipping is certainly preferred to killing the animal. Attention should be paid to this issue, however, and, where appropriate, survival estimates should be undertaken to determine the health implications of this procedure. Also, antiseptic and analgesic protocols can be considered to ensure that wounds where tissue samples are excised are at low risk of secondary infection (Chevalier *et al*. 2017).

In summary, modification of Longcore’s original *Bd*-isolation protocol (Longcore *et al*. 1999) has enabled a broad community of scientists to engage with research on emerging chytrid pathogens of amphibians. This research has had an impact worldwide, and is contributing to the ongoing dialogue that is occurring between scientists, conservationists and policy-makers about how we might mitigate against these infections now and into the future.

## Data accessibility

https://five.epicollect.net/project/bd-global-isolation-protocol

## References

1. Aanensen, D. M., D. M. Huntley, E. J. Feil, F. al-Own and B. G. Spratt (2009). "EpiCollect: Linking Smartphones to Web Applications for Epidemiology, Ecology and Community Data Collection." Plos One 4(9).

2. Aanensen, D. M., D. M. Huntley, M. Menegazzo, C. I. Powell and B. G. Spratt (2014). "EpiCollect+: linking smartphones to web applications for complex data collection projects." F1000Res 3: 199.

3. Barr, D. J. S. (1987). Isolation, culture and identification of Chytridiales, Spizellomycetales, and Hypocreales. Zoosporic fungi in teaching and research. M. S. Fuller and A. Jaworski. Athens, Georgia, Southeastern Publishing Corp.: p 118–120.

4. Bataille, A., J. J. Fong, M. Cha, G. O. U. Wogan, H. J. Baek, H. Lee, M. S. Min and B. Waldman (2013). "Genetic evidence for a high diversity and wide distribution of endemic strains of the pathogenic chytrid fungus Batrachochytrium dendrobatidis in wild Asian amphibians." Molecular Ecology 22(16): 4196–4209.

5. Becker, C. G., S. E. Greenspan, K. E. Tracy, J. A. Dash, C. Lambertini, T. S. Jenkinson, D. S. Leite, L. F. Toledo, J. E. Longcore, T. Y. James and K. R. Zamudio (2017). "Variation in phenotype and virulence among enzootic and panzootic amphibian chytrid lineages." Fungal Ecology 26: 45–50.

6. Berger, L., R. Speare, P. Daszak, D. E. Green, A. A. Cunningham, C. L. Goggin, R. Slocombe, M. A. Ragan, A. H. Hyatt, K. R. McDonald, H. B. Hines, K. R. Lips, G. Marantelli and H. Parkes (1998). "Chytridiomycosis causes amphibian mortality associated with population declines in the rain forests of Australia and Central America." Proceedings of the National Academy of Science, USA 95: 9031–9036.

7. Blooi, M., A. Martel, F. Haesebrouck, F. Vercammen, D. Bonte and F. Pasmans (2015). "Treatment of urodelans based on temperature dependent infection dynamics of Batrachochytrium salamandrivorans." Scientific Reports 5.

8. Blooi, M., F. Pasmans, J. E. Longcore, A. Spitzen-van der Sluijs, F. Vercammen and A. Martel (2013). "Duplex Real-Time PCR for Rapid Simultaneous Detection of Batrachochytrium dendrobatidis and Batrachochytrium salamandrivorans in Amphibian Samples." Journal of Clinical Microbiology 51(12): 4173–4177.

9. Boyle, D. G., D. B. Boyle, V. Olsen, J. A. T. Morgan and A. D. Hyatt (2004). "Rapid quantitative detection of chytridiomycosis (Batrachochytrium dendrobatidis) in amphibian samples using real-time Taqman PCR assay." Diseases of Aquatic Organisms 60(2): 141–148.

10. Boyle, D. G., A. D. Hyatt, P. Daszak, L. Berger, J. E. Longcore, D. Porter, S. G. Hengstberger and V. Olsen (2003). "Cryo-archiving of Batrachochytrium dendrobatidis and other chytridiomycetes." Dis Aquat Organ 56(1): 59–64.

11. Canessa, S., C. Bozzuto, E. H. C. Grant, S. S. Cruickshank, M. C. Fisher, J. C. Koella, S. Lötters, A. Martel, F. Pasmans, B. C. Scheele, A. Spitzen-van der Sluijs, S. S. and B. R. Schmidt (2018). "Decision making for mitigating emerging wildlife diseases: from theory to practice for an emerging fungal pathogen of salamanders..” Journal of Applied Ecology.

12. Chevalier, H., O. Calvez, A. MArtinez-Silvestre, D. Picard, S. Guerin, F. Isselin-Nondedeu, A. Riberon and A. Trochet (2017). "Marking techniques in the Marbled Newt (Triturus marmoratus): PIT-Tag and tracking device implant protocols." Acta Herpetologica 12(1): 79–88.

13. Dillon, M. J., A. E. Bowkett, M. J. Bungard, K. M. Beckman, M. F. O’Brien, K. Bates, M. C. Fisher, J. R. Stevens and C. R. Thornton (2017). "Tracking the amphibian pathogens Batrachochytrium dendrobatidis and Batrachochytrium salamandrivorans using a highly specific monoclonal antibody and lateral- flow technology." Microbial Biotechnology 10(2): 381–394.

14. Farrer, R. A. and M. C. Fisher (2017). "Describing Genomic and Epigenomic Traits Underpinning Emerging Fungal Pathogens." Adv Genet 100: 73–140.

15. Farrer, R. A., D. A. Henk, T. W. J. Garner, F. Balloux, D. C. Woodhams and M. C. Fisher (2013). "Chromosomal Copy Number Variation, Selection and Uneven Rates of Recombination Reveal Cryptic Genome Diversity Linked to Pathogenicity." Plos Genetics 9(8).

16. Farrer, R. A., A. Martel, E. Verbrugghe, A. Abouelleil, R. Ducatelle, J. E. Longcore, T. Y. James, F. Pasmans, M. C. Fisher and C. A. Cuomo (2017). "Genomic innovations linked to infection strategies across emerging pathogenic chytrid fungi." Nat Commun 8: 14742.

17. Farrer, R. A., L. A. Weinert, J. Bielby, T. W. J. Garner, F. Balloux, F. Clare, J. Bosch, A. A. Cunningham, C. Weldon, L. H. du Preez, L. Anderson, S. L. K. Pond, R. Shahar-Golan, D. A. Henk and M. C. Fisher (2011). "Multiple emergences of genetically diverse amphibian-infecting chytrids include a globalized hypervirulent recombinant lineage." Proceedings of the National Academy of Sciences of the United States of America 108(46): 18732–18736.

18. Fellers, G. M., D. E. Green and J. E. Longcore (2001). "Oral chytridiomycosis in the mountain yellow-legged frog (Rana muscosa)." Copeia(4): 945–953.

19. Fisher, M. C., J. Bosch, Z. Yin, D. A. Stead, J. Walker, L. Selway, A. J. P. Brown, L. A. Walker, N. A. R. Gow, J. E. Stajich and T. W. J. Garner (2009). "Proteomic and phenotypic profiling of the amphibian pathogen Batrachochytrium dendrobatidis shows that genotype is linked to virulence." Molecular Ecology 18(3): 415–429.

20. Fisher, M. C. and T. W. J. Garner (2007). "The relationship between the introduction of Batrachochytrium dendrobatidis, the international trade in amphibians and introduced amphibian species." Fungal Biology Reviews 21: 2–9.

21. Fisher, M. C., T. W. J. Garner and S. F. Walker (2009). "Global Emergence of Batrachochytrium dendrobatidis and Amphibian Chytridiomycosis in Space, Time, and Host." Annual Review of Microbiology 63: 291–310.

22. Fisher, M. C., D. A. Henk, C. J. Briggs, J. S. Brownstein, L. C. Madoff, S. L. McCraw and S. J. Gurr (2012). "Emerging fungal threats to animal, plant and ecosystem health." Nature 484(7393): 186–194.

23. Fisher, M. C., B. R. Schmidt, K. Henle, D. S. Schmeller, J. Bosch, D. M. Aanensen and T. J. W. Garner (2012). "RACE: Risk Assessment of Chytridiomycosis to European Amphibian Biodiversity.." Froglog(101): 45–47.

24. Funk, W. C., M. A. Donnelly and K. R. Lips (2005). "Alternative views of amphibian toe-clipping." Nature 433(7023): 193–193.

25. Garner, T. W., B. R. Schmidt, A. Martel, F. Pasmans, E. Muths, A. A. Cunningham, C. Weldon, M. C. Fisher and J. Bosch (2016). "Mitigating amphibian chytridiomycoses in nature." Philos Trans R Soc Lond B Biol Sci 371(1709).

26. Garner, T. W., S. Walker, J. Bosch, S. Leech, M. Rowcliffe, A. A. Cunningham and M. C. Fisher (2009). "Life history trade-offs influence mortality associated with the amphibian pathogen Batrachochytrium dendrobatidis." Oikos(118): 783–791.

27. Gentz, E. J. (2007). "Medicine and surgery of amphibians." ILAR J (48): 255–259.

28. James, T. Y., A. Litvintseva, R. Vilgalys, J. A. Morgan, J. W. Taylor, M. C. Fisher, L. Berger, C. Weldon, L. H. du Preez and J. Longcore (2009). "Rapid expansion of an emerging fungal disease into declining and healthy amphibian populations." PLoS Pathogens 5(5): e1000458.

29. James, T. Y., L. F. Toledo, D. Rodder, D. D. Leite, A. M. Belasen, C. M. Betancourt-Roman, T. S. Jenkinson, C. Soto-Azat, C. Lambertini, A. V. Longo, J. Ruggeri, J. P. Collins, P. A. Burrowes, K. R. Lips, K. R. Zamudio and J. E. Longcore (2015). "Disentangling host, pathogen, and environmental determinants of a recently emerged wildlife disease: lessons from the first 15years of amphibian chytridiomycosis research." Ecology and Evolution 5(18): 4079–4097.

30. Jenkinson, T. S., C. M. B. Roman, C. Lambertini, A. Valencia-Aguilar, D. Rodriguez, C. H. L. Nunes-de-Almeida, J. Ruggeri, A. M. Belasen, D. D. Leite, K. R. Zamudio, J. E. Longcore, L. F. Toledo and T. Y. James (2016). "Amphibian-killing chytrid in Brazil comprises both locally endemic and globally expanding populations." Molecular Ecology 25(13): 2978–2996.

31. Laking, A. E., H. N. Ngo, F. Pasmans, A. Martel and T. T. Nguyen (2017). "Batrachochytrium salamandrivorans is the predominant chytrid fungus in Vietnamese salamanders." Scientific Reports 7.

32. Longcore, J. E., A. P. Pessier and D. K. Nichols (1999). "Batrachochytrium dendrobatidis gen et sp nov, a chytrid pathogenic to amphibians." Mycologia 91(2): 219–227.

33. Martel, A., A. Spitzen-van der Sluijs, M. Blooi, W. Bert, R. Ducatelle, M. C. Fisher, A. Woeltjes, W. Bosman, K. Chiers, F. Bossuyt and F. Pasmans (2013). "Batrachochytrium salamandrivorans sp nov causes lethal chytridiomycosis in amphibians." Proceedings of the National Academy of Sciences of the United States of America 110(38): 15325–15329.

34. May, R. M. (2004). "Ecology - Ethics and amphibians." Nature 431(7007): 403–403.

35. McCarthy, M. A. and K. M. Parris (2004). "Clarifying the effect of toe clipping on frogs with Bayesian statistics." Journal of Applied Ecology 41(4): 780–786.

36. Mendelson, J. R., 3rd, K. R. Lips, R. W. Gagliardo, G. B. Rabb, J. P. Collins, J. E. Diffendorfer, P. Daszak, D. R. Ibanez, K. C. Zippel, D. P. Lawson, K. M. Wright, S. N. Stuart, C. Gascon, H. R. da Silva, P. A. Burrowes, R. L. Joglar, E. La Marca, S. Lotters, L. H. du Preez, C. Weldon, A. Hyatt, J. V. Rodriguez-Mahecha, S. Hunt, H. Robertson, B. Lock, C. J. Raxworthy, D. R. Frost, R. C. Lacy, R. A. Alford, J. A. Campbell, G. Parra-Olea, F. Bolanos, J. J. Domingo, T. Halliday, J. B. Murphy, M. H. Wake, L. A. Coloma, S. L. Kuzmin, M. S. Price, K. M. Howell, M. Lau, R. Pethiyagoda, M. Boone, M. J. Lannoo, A. R. Blaustein, A. Dobson, R. A. Griffiths, M. L. Crump, D. B. Wake and E. D. Brodie, Jr. (2006). "Biodiversity. Confronting amphibian declines and extinctions." Science 313(5783): 48.

37. Ott, J. A. and D. E. Scott (1999). "Effects of toe-clipping and PIT-tagging on growth and survival in metamorphic Ambystoma opacum." Journal of Herpetology 33(2): 344–348.

38. Piotrowski, J. S., S. L. Annis and J. E. Longcore (2004). "Physiology of Batrachochytrium dendrobatidis, a chytrid pathogen of amphibians." Mycologia 96(1): 9–15.

39. Ribas, L., M. S. Li, B. J. Doddington, J. Robert, J. A. Seidel, J. S. Kroll, L. B. Zimmerman, N. C. Grassly, T. W. J. Garner and M. C. Fisher (2009). "Expression Profiling the Temperature-Dependent Amphibian Response to Infection by Batrachochytrium dendrobatidis." PLoS One 4(12): -.

40. Rosenblum, E. B., T. J. Poorten, S. Joneson and M. Settles (2012). "Substrate-Specific Gene Expression in Batrachochytrium dendrobatidis, the Chytrid Pathogen of Amphibians." Plos One 7(11).

41. Rosenblum, E. B., T. J. Poorten, M. Settles and G. K. Murdoch (2012). "Only skin deep: shared genetic response to the deadly chytrid fungus in susceptible frog species." Molecular Ecology 21(13): 3110–3120.

42. Schloegel, L. M., C. M. Ferreira, T. Y. James, M. Hipolito, J. E. Longcore, A. D. Hyatt, M. Yabsley, A. M. C. R. P. F. Martins, R. Mazzoni, A. J. Davies and P. Daszak (2010). "The North American bullfrog as a reservoir for the spread of Batrachochytrium dendrobatidis in Brazil." Animal Conservation 13: 53–61.

43. Schloegel, L. M., A. M. Picco, A. M. Kilpatrick, A. J. Davies, A. D. Hyatt and P. Daszak (2009). "Magnitude of the US trade in amphibians and presence of Batrachochytrium dendrobatidis and ranavirus infection in imported North American bullfrogs (Rana catesbeiana)." Biological Conservation 142(7): 1420–1426.

44. Schloegel, L. M., L. F. Toledo, J. E. Longcore, S. E. Greenspan, C. A. Vieira, M. Lee, S. Zhao, C. Wangen, C. M. Ferreira, M. Hipolito, A. J. Davies, C. A. Cuomo, P. Daszak and T. Y. James (2012). "Novel, panzootic and hybrid genotypes of amphibian chytridiomycosis associated with the bullfrog trade." Molecular Ecology 21(21): 5162–5177.

45. Schmeller, D. S., M. Blooi, A. Martel, T. W. J. Garner, M. C. Fisher, F. Azemar, F. C. Clare, C. Leclerc, L. Jager, M. Guevara-Nieto, A. Loyau and F. Pasmans (2014). "Microscopic Aquatic Predators Strongly Affect Infection Dynamics of a Globally Emerged Pathogen." Current Biology 24(2): 176–180.

46. Schoch, C. L., K. A. Seifert, S. Huhndorf, V. Robert, J. L. Spouge, C. A. Levesque, W. Chen, E. Bolchacova, K. Voigt, P. W. Crous, A. N. Miller, M. J. Wingfield, M. C. Aime, K. D. An, F. Y. Bai, R. W. Barreto, D. Begerow, M. J. Bergeron, M. Blackwell, T. Boekhout, M. Bogale, N. Boonyuen, A. R. Burgaz, B. Buyck, L. Cai, Q. Cai, G. Cardinali, P. Chaverri, B. J. Coppins, A. Crespo, P. Cubas, C. Cummings, U. Damm, Z. W. de Beer, G. S. de Hoog, R. Del-Prado, D. B, J. Dieguez-Uribeondo, P. K. Divakar, B. Douglas, M. Duenas, T. A. Duong, U. Eberhardt, J. E. Edwards, M. S. Elshahed, K. Fliegerova, M. Furtado, M. A. Garcia, Z. W. Ge, G. W. Griffith, K. Griffiths, J. Z. Groenewald, M. Groenewald, M. Grube, M. Gryzenhout, L. D. Guo, F. Hagen, S. Hambleton, R. C. Hamelin, K. Hansen, P. Harrold, G. Heller, G. Herrera, K. Hirayama, Y. Hirooka, H. M. Ho, K. Hoffmann, V. Hofstetter, F. Hognabba, P. M. Hollingsworth, S. B. Hong, K. Hosaka, J. Houbraken, K. Hughes, S. Huhtinen, K. D. Hyde, T. James, E. M. Johnson, J. E. Johnson, P. R. Johnston, E. B. Jones, L. J. Kelly, P. M. Kirk, D. G. Knapp, U. Koljalg, K. Gm, C. P. Kurtzman, S. Landvik, S. D. Leavitt, A. S. Liggenstoffer, K. Liimatainen, L. Lombard, J. J. Luangsa-Ard, H. T. Lumbsch, H. Maganti, S. S. Maharachchikumbura, M. P. Martin, T. W. May, A. R. McTaggart, A. S. Methven, W. Meyer, J. M. Moncalvo, S. Mongkolsamrit, L. G. Nagy, R. H. Nilsson, T. Niskanen, I. Nyilasi, G. Okada, I. Okane, I. Olariaga, J. Otte, T. Papp, D. Park, T. Petkovits, R. Pino-Bodas, W. Quaedvlieg, H. A. Raja, D. Redecker, R. T, C. Ruibal, J. M. Sarmiento-Ramirez, I. Schmitt, A. Schussler, C. Shearer, K. Sotome, F. O. Stefani, S. Stenroos, B. Stielow, H. Stockinger, S. Suetrong, S. O. Suh, G. H. Sung, M. Suzuki, K. Tanaka, L. Tedersoo, M. T. Telleria, E. Tretter, W. A. Untereiner, H. Urbina, C. Vagvolgyi, A. Vialle, T. D. Vu, G. Walther, Q. M. Wang, Y. Wang, B. S. Weir, M. Weiss, M. M. White, J. Xu, R. Yahr, Z. L. Yang, A. Yurkov, J. C. Zamora, N. Zhang, W. Y. Zhuang, D. Schindel and F. B. Consortium (2012). "Nuclear ribosomal internal transcribed spacer (ITS) region as a universal DNA barcode marker for Fungi." Proceedings of the National Academy of Sciences of the United States of America109(16): 6241–6246.

47. Skerratt, L. F., L. Berger, H. B. Hines, K. R. McDonald, D. Mendez and R. Speare (2008). "Survey protocol for detecting chytridiomycosis in all Australian frog populations." Diseases of Aquatic Organisms 80(2): 85–94.

48. Smith, K. G. and C. Weldon (2007). "A Conceptual Framework for Detecting Oral Chytridiomycosis in Tadpoles." Copeia 4: 1024–1028.

49. Stuart, S. N., J. S. Chanson, N. A. Cox, B. E. Young, A. S. Rodrigues, D. L. Fischman and R. W. Waller (2004). "Status and trends of amphibian declines and extinctions worldwide." Science 306(5702): 1783–1786.

50. Swei, A., J. J. L. Rowley, D. Rodder, M. L. L. Diesmos, A. C. Diesmos, C. J. Briggs, R. Brown, T. T. Cao, T. L. Cheng, R. A. Chong, B. Han, J. M. Hero, H. D. Hoang, M. D. Kusrini, T. T. L. Duong, J. A. McGuire, M. Meegaskumbura, M. S. Min, D. G. Mulcahy, T. Neang, S. Phimmachak, D. Q. Rao, N. M. Reeder, S. D. Schoville, N. Sivongxay, N. Srei, M. Stock, B. L. Stuart, L. S. Torres, T. A. T. Dao, T. S. Tunstall, D. Vieites and V. T. Vredenburg (2011). "Is Chytridiomycosis an Emerging Infectious Disease in Asia?" Plos One 6(8).

51. Torreilles, S. L., D. E. McClure and S. L. Green (2009). "Evaluation and Refinement of Euthanasia Methods for Xenopus laevis." Journal of the American Association for Laboratory Animal Science 48(5): 512–516.

52. Ulmar Grafe, T., M. M. Stewart, K. P. Lampert and M.-O. Rödel (2011). "Putting Toe Clipping into Perspective: A Viable Method for Marking Anurans." Journal of Herpetology 45(1): 28–35.

53. Voyles, J., L. Berger, S. Young, R. Speare, R. Webb, J. Warner, D. Rudd, R. Campbell and L. F. Skerratt (2007). "Electrolyte depletion and osmotic imbalance in amphibians with chytridiomycosis." Diseases of Aquatic Organisms 77(2): 113–118.

54. Walker, S., J. Bosch, T. Y. James, A. Litvintseva, J. A. O. Valls, S. Pina, G. Garcia, G. Abadie-Rosa, A. A. Cunningham, S. Hole, R. Griffiths and M. C. Fisher (2008). "Invasive pathogens threaten species recovery programs." Current Biology 18: R853–R854.

55. Webb, R., L. Berger, D. Mendez and R. Speare (2005). "MS-222 (tricaine methane sulfonate) does not kill the amphibian chytrid fungus Batrachochytrium dendrobatidis." Diseases of Aquatic Organisms 68(1): 89–90.

